# Functionalizing hydrogel nanovials with vesicles mimicking antigen-presenting vesicles and cancer exosomes improves T cell capture and activation

**DOI:** 10.1101/2025.06.24.661128

**Authors:** Blade A. Olson, Michael Mellody, Citradewi Soemardy, Zhiyuan Mao, Anna Mei, Katie Lippert, Magnus A. G. Hoffmann, Dino Di Carlo, Stephen L. Mayo

## Abstract

Recent advances have demonstrated the application of microcavity-containing hydrogel microparticles, known as nanovials, for the massively parallel and high-throughput screening of therapeutic T cell populations for adoptive cell therapies. Nanovial cavities coated with peptide-MHC (pMHC) or antigen tetramers selectively bind to their cognate T cell receptor (TCR) or chimeric antigen receptor (CAR) to activate T cells and capture secreted cytokines. However, binding of tetramers or recombinantly expressed antigen by T cells is not always correlated with T cell activation or cytotoxicity as the binding interface is not fully representative of the natural immunological synapse formed between T cells and professional antigen-presenting cells (APCs). Here, we leverage the recent discovery of an ESCRT- and ALIX-binding region (EABR) sequence to generate antigen-presenting vesicles and cancer-mimicking exosomes from standard HEK293T and Expi293F cell cultures. EABR-mediated vesicles present natural, full-length oncologically-relevant membrane proteins embedded in lipid bilayers to functionalize the nanovial cavity with cell-like membranes. These hydrogel nanovials functionalized with the EABR-mediated vesicles show improved T cell capture of 1G4 T cells and enhanced activation of HER2 CAR-T cells compared to hydrogel surfaces functionalized with recombinantly-expressed soluble proteins.

## Introduction

Adoptive cell therapy has shown remarkable effectiveness in the treatment of advanced hematological malignancies, and chimeric antigen receptor (CAR) T cell therapy has achieved high rates of complete recovery in patients with treatment-resistant, relapsed or refractory B cell malignancies^1–6^. However, a major hurdle in developing effective immunotherapies is the identification of reactive TCRs or CARs that can recognize targets with sufficient affinity, selectivity, and potency^7^. Further, the T cells that are identified to harbor a reactive TCR of sufficient affinity are often limited by suboptimal *ex vivo* T cell expansion rates and final T cell products ultimately have dysfunctional or limited clinical functionality^8, 9^. Current tools for enriching and screening promising T cell populations typically rely on TCR or CAR affinity by staining with peptide-MHC (pMHC) or antigen tetramers. However, binding of multimers to T cells is not always correlated with T cell activation or cytotoxicity^10^ as the context in which these pMHC signals are presented is understandably not representative of how they are naturally presented by professional antigen-presenting cells (APCs) or cancer cells.^11^

To efficiently screen through a pool of T cells and identify those with therapeutic potential, Koo and Mao et al. recently reported a high-throughput approach to selectively capture single antigen-specific T cells in cavity-containing hydrogel microparticles^12^, called nanovials.^13^ Each nanovial acts as both an artificial APC that presents pMHC molecules to activate cells with cognate TCRs, and as a capture site for secreted effector molecules, such as IFN-γ and granzyme B, that enable high-throughput sorting of therapeutically relevant T cells by flow cytometry. While this capture and activation method was successful in isolating previously unidentified reactive T cells, we hypothesized that improved selection and activation would be possible by incorporating the immune-activating signaling molecules into lipid membranes with high fluidity. Signaling molecules in lipid membranes with high fluidity have been shown to activate T cells better than membranes with low fluidity^14^ due to the improved ability to facilitate receptor clustering in the supramolecular activation complex, or “immunological synapse”, that forms between an APC and a T cell during antigen presentation and subsequent immune activation^11^.

To recreate the potential benefits of these physiological lipid bilayers, researchers have previously imbued bioinert materials with bioactive cell-like membranes through extrusion, sonication, or centrifugation, which typically involve coating with non-specific cell lysates and may be difficult to achieve with heterogenous surfaces^15^. Recently, synthetic liposomes were coated on mesoporous silica micro-rods^16^ to generate supported lipid bilayers harboring combinations of biotinylated T cell activation cues, such as pMHC and αCD28 antibodies, for the rapid expansion of highly functional T cells; while this method enabled biotinylated signaling molecules to have membrane fluidity for improved T cell activation, a known limitation of this approach is that many membrane proteins may be insoluble or display altered or absent activities if synthesized recombinantly outside of a lipid bilayer^17, 18^; thus, an ideal functionalization strategy would allow for natural, full-length cell membrane proteins embedded in highly fluid lipid bilayers to coat the surface of the intended bioactive material without the need for recombinant expression of a soluble protein for surface conjugation. Accordingly, extracellular vesicles featuring natural, full-length membrane proteins have been increasingly pursued as a coating for biomaterials with successes seen in both tissue engineering^19–21^ and immunomodulatory^22–25^ settings. However, an ongoing struggle with natural extracellular vesicles is that their low expression yield^26^ and the heterogeneity of their protein composition^27^ precludes reliable experimentation and manufacturing. The surface density of certain immunoregulatory proteins, for example, is a critical parameter for immune activation^28^, as previous studies have demonstrated that exosomes or cell sonicates must be derived from APCs that express a high density of pMHCI and CD80/CD54 complexes to induce T cell responses^29^.

Here, we demonstrate the expression and purification of high concentrations of extracellular vesicles that are densely coated with desired compositions of immunoregulatory membrane proteins, which we then use to functionalize the surfaces of nanovial cavities for improved T cell capture and activation. As initially described by Hoffmann et al., vesicles are generated by transfecting standard HEK293T and Expi293F cell cultures with protein constructs containing an appended ESCRT- and ALIX-binding region (EABR) sequence^30^ to the cytoplasmic tail of the full-length membrane protein; this EABR sequence recruits ESCRT proteins to induce the budding of extracellular vesicles that can optionally mimic cancer-derived HER2-presenting extracellular vesicles or antigen-presenting vesicles (APVs)^31^ co-presenting both pMHCI and the immunoregulatory proteins CD274, CD83, or CD80. As a demonstration of the biological relevance of these engineered extracellular vesicles for advancing adoptive cell therapies, we show the facile coating of heterogeneous surfaces of microparticle hydrogel nanovials with these engineered extracellular vesicles to improve the loading and activation of therapeutically-relevant T cells when compared to their soluble recombinantly expressed variants. T cells captured in these nanovials can be isolated for further downstream analysis, enabling both improved high-throughput single cell discovery of T cells with therapeutic potential as well as future single cell studies of extracellular vesicle membrane protein composition on T cell activation and differentiation.

## Results

### Hydrogel nanovials enable single-cell comparison of T cell activation by extracellular vesicles

Recent work by Hoffmann et al. demonstrated a generalizable method for inducing the release of transfected membrane proteins on densely coated, cell-derived extracellular vesicles^30^. Appending an endosomal sorting complex required for transport (ESCRT)- and ALG-2-interacting protein X (ALIX)- binding region (EABR) to the cytoplasmic tail of membrane proteins directly recruits the ESCRT machinery and induces the release of vesicles displaying the EABR-tagged protein (Figure 1A). Transfecting common HEK293T or Expi293F cell cultures with DNA plasmids encoding EABR-tagged membrane proteins generates vesicles that can be purified from the cell culture supernatant by ultracentrifugation on a 20% sucrose gradient as is typical for the purification of viral particles and exosomes^32^ (Figure 1B). We hypothesized that these cell-derived extracellular vesicles would activate T cells more strongly than soluble protein by presenting cancer antigens and pMHCI complexes in a more physiologically-relevant format.

**Figure 1.**
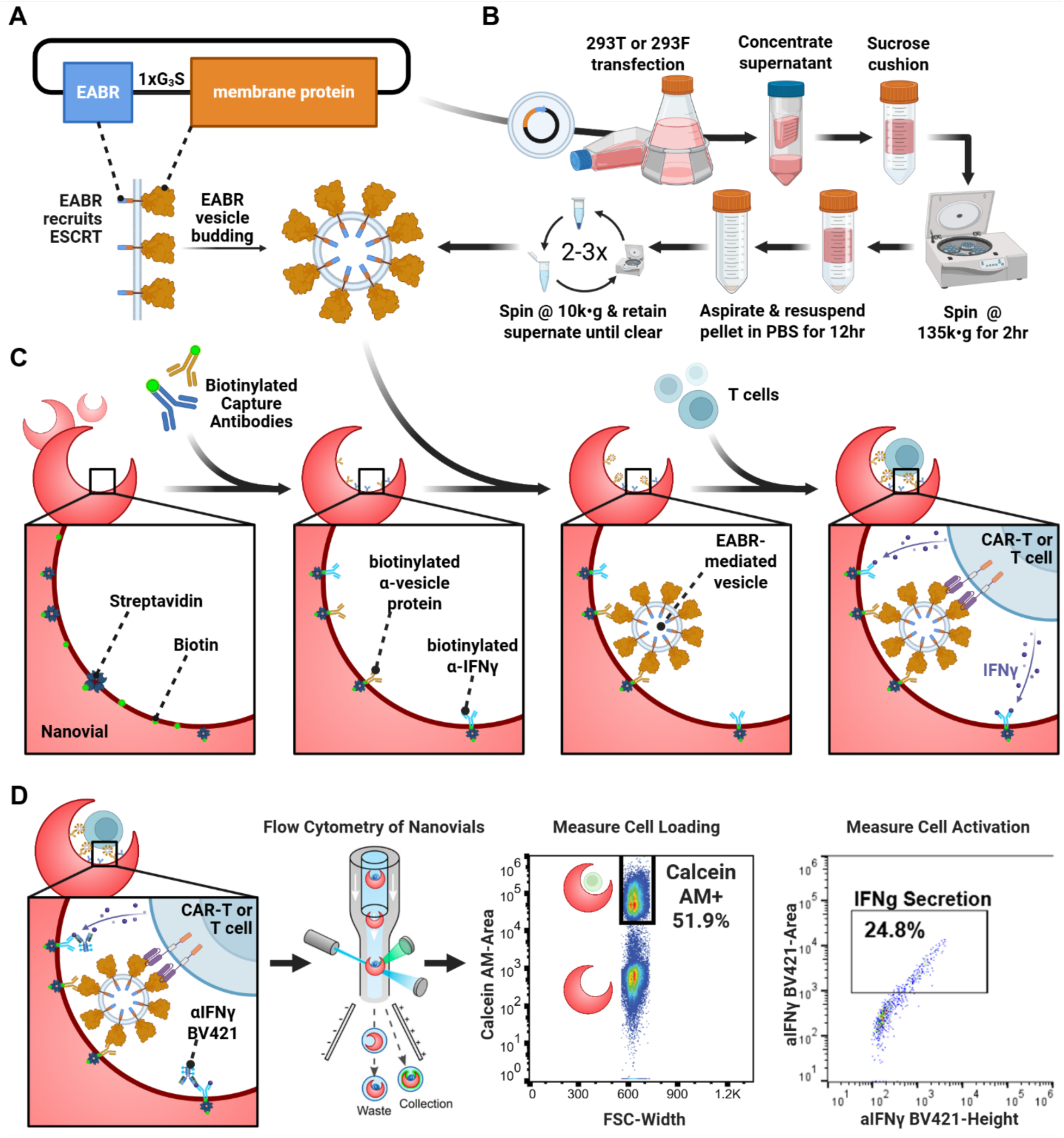
Engineered extracellular vesicles improve single cell capture and T cell activation of hydrogel nanovials. (A) Top: schematic of a plasmid construct containing a membrane protein fused to a Gly_3_Ser spacer, and an EABR vesicle-forming sequence. Bottom: schematic of membrane-bound protein on a cell surface containing cytoplasmic tail EABR additions that induce budding of a vesicle comprising a lipid bilayer with embedded membrane proteins that contain the EABR sequence. (B) Schematic showing production and purification of the engineered EABR-mediated extracellular vesicles. (C) Post-fabrication biotinylation of hydrogel nanovials via gelatin are then functionalized with streptavidin, followed by biotinylated capture antibodies. Capture antibodies can be any protein present on the vesicle surface, including the EABR-tagged membrane protein that induces vesicle formation. Cells, such as CAR-T cells, bind to the captured extracellular vesicles within the nanovial cavity, and cell activation cytokines are captured using the biotinylated capture antbiodies covering the nanovial surface. (D) Fluorescent sandwich antibodies detect cytokines released by activated cells that are captured within the nanovial to enable single-cell measurements of both cell capture and cell activation by flow cytometry.

To evaluate the performance of these engineered extracellular vesicles for cell capture and activation, structured hydrogel microparticles, or “nanovials”^13^, were manufactured to hold single cells in inner cavities functionalized with the EABR-mediated extracellular vesicles (Figure 1C). Briefly, a microfluidic device generates uniform water-in-oil emulsions, where aqueous two-phase separation of polyethylene glycol (PEG) and gelatin occurs to create millions of monodisperse PEG-based nanovials with an inner cavity selectively coated with biotinylated gelatin. Multiple biotinylated antibodies or peptide-MHC (pMHC) monomers with epitopes of interest can be flexibly linked to the cavity’s interior through streptavidin-biotin noncovalent interactions^12^. These antibodies can be used for non-specific T cell capture (e.g., CD45), for capture of T cell activation cytokines (e.g., IFN-γ), or for capture of EABR-mediated engineered extracellular vesicles by binding to transfected membrane proteins present on the surface of the vesicles (e.g., HER2, CD54, CD274, CD83, CD80, or pMHCI). CAR-T or T cells are captured in the cavity of nanovials by binding to the EABR-tagged membrane proteins of the extracellular vesicles functionalizing the nanovial surface. Subsequent T cell activation and IFN-γ release is measured by a sandwich antibody for IFN-γ captured in the nanovial cavity’s interior (Figure 1D). Extracellular vesicles, T cells, and their associated secretions are then analyzed and sorted using high-throughput commercial flow cytometers, which can conveniently isolate therapeutically-relevant, antigen-reactive T cells. This massively parallel and high-throughput system provides a single-cell resolution of T cell activation to compare the performance of T cell capture and activation by extracellular vesicles versus recombinantly expressed pMHCI and cancer antigen.

### Co-expression of SCT pMHCI/EABR and CD274, CD83, or CD80 generates immunoregulatory antigen-presenting vesicles

We previously demonstrated the engineered assembly and budding of pMHCI-displaying APVs from the cell surface of non-immune cells by DNA transfection of an engineered single-chain heterotrimer (SCT) pMHCI^31^. Expanding on this work, we simultaneously transfected cell cultures with two constructs encoding a single-chain heterotrimer peptide:MHCI (SCT pMHCI) complex and either programmed cell death ligand 1 (PD-L1) CD274, immunoregulatory CD83, or the costimulatory signal CD80, each of which has been found previously on immune cell surfaces^33^, immunoregulatory APVs^34^, and cancer-derived exosomes^35, 36^ (Figure 2A). Continuing from our earlier work, we utilized the “D9” variant of SCT pMHCI^37–39^, containing Y84C and A139C substitutions in the alpha heavy chain that were previously reported to improve both thermostability and TCR binding.

**Figure 2.**
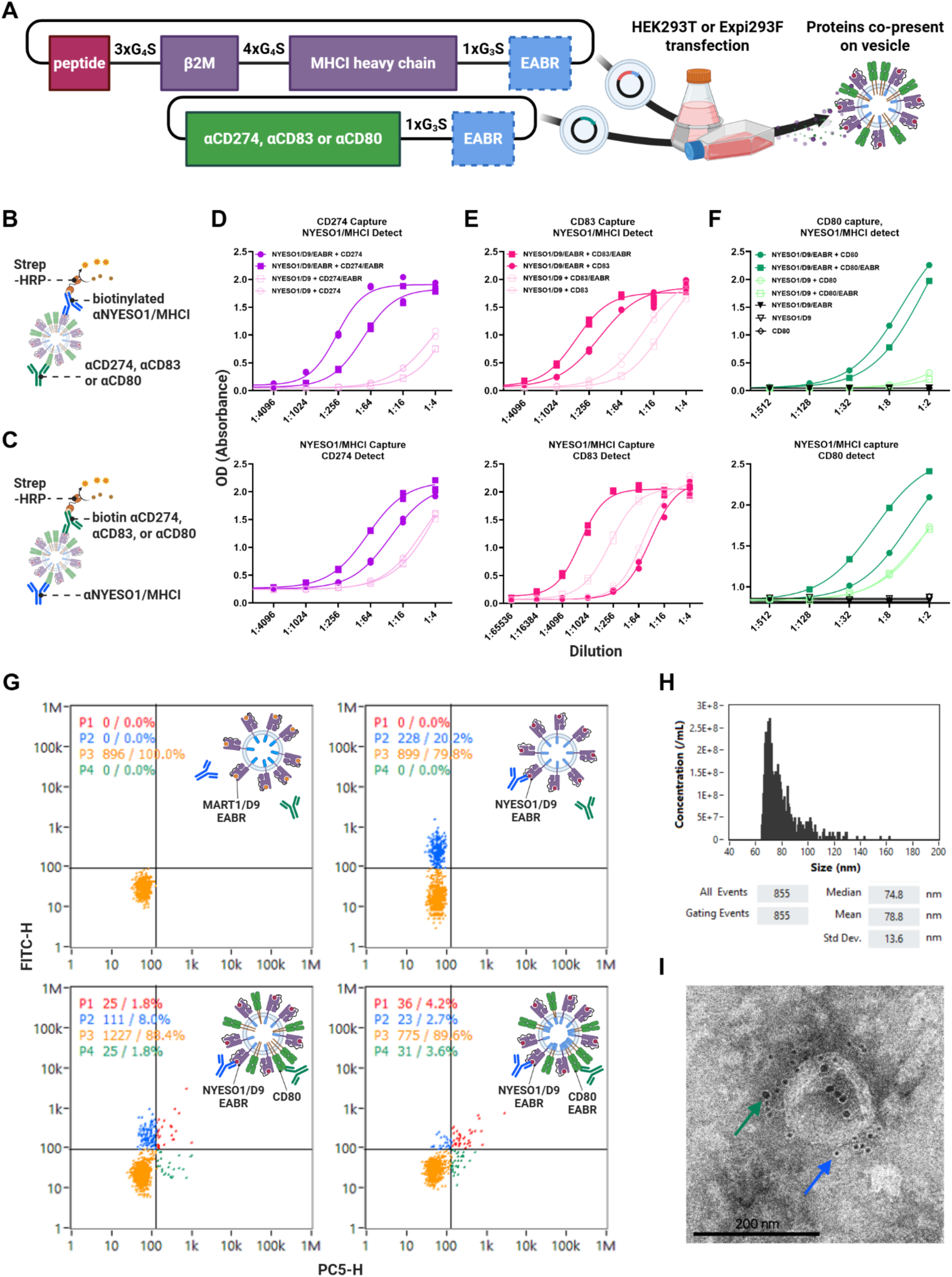
Co-expression of SCT pMHCI/EABR and CD274, CD83 or CD80 generates immunoregulatory antigen-presenting vesicles. (A) Schematic showing the plasmid constructs for vesicles co-presenting NYESO1/HLA-A*0201 and either CD274, CD83, or CD80 with an “EABR” vesicle-forming sequence optionally appended to the cytoplasmic tail. The plasmid is transfected into HEK293T or Expi293F cells and vesicles are harvested as shown previously in Figure 1B. (B) Schematic of the assay performed for the top row of Fig. (3C-E). Expression of immunoregulatory membrane proteins CD274, CD83, or CD80 on engineered extracellular vesicles were captured with plated antibody, then co-presented NYESO1/HLA-A*02:01 were detected and quantified with biotinylated anti-NYESO:HLA-A*02:01 and HRP-conjugated streptavidin (strep-HRP). (C) Schematic of the assay performed for the bottom row of Fig. C-E. Identical assay to (B), but the capture antibody and detection antibody have been swapped. (D) Sandwich ELISA of APVs purified by ultracentrifugation from transfected Expi293F cells using anti-CD274 for capture, biotinylated anti-NYESO1:HLA-A*02:01 (3M4E5) for detection, and HRP-conjugated streptavidin secondary. Cells were co-transfected with SCT NYESO1/D9/EABR and CD274, SCT NYESO1/D9/EABR and CD274/EABR, SCT NYESO1/D9 and CD274/EABR, or SCT NYESO1/D9 and CD274 constructs (2 replicates per dilution). (E) Sandwich ELISA of APVs purified by ultracentrifugation from transfected Expi293F cells using anti-CD83 for capture, biotinylated anti-NYESO1:HLA-A*02:01 (3M4E5) for detection, and HRP-conjugated streptavidin secondary. Cells were co-transfected with SCT NYESO1/D9/EABR and CD83, SCT NYESO1/D9/EABR and CD83/EABR, SCT NYESO1/D9 and CD83/EABR, or SCT NYESO1/D9 and CD83 constructs (2 replicates per dilution). (F) Sandwich ELISA of APVs purified by ultracentrifugation from transfected HEK293T cells using anti-CD80 for capture, biotinylated anti-NYESO1:HLA-A*02:01 (3M4E5) for detection, and HRP-conjugated streptavidin secondary. Cells were co-transfected with SCT NYESO1/D9/EABR and CD80, SCT NYESO1/D9/EABR and CD80/EABR, SCT NYESO1/D9 and CD80/EABR, or SCTNYESO1/D9 and CD80 constructs (2 replicates per dilution).(G) Nano flow cytometry (NanoFCM) of APVs purified by ultracentrifugation from transfected HEK293T cells, and co-stained with APC anti-CD80 and Alexa488 anti-NYESO1:HLA-A*02:01 (3M4E5). Top-Left: Vesicles purified from HEK293T transfected with the MART1/D9/EABR construct were used for gating non-specific signal from irrelevant extracellular vesicles. Top-Right: Vesicles from HEK293T cells transfected with the NYESO1/D9/EABR construct. Bottom-Left: Vesicles from HEK293T cells transfected with NYESO1/D9/EABR and CD80 constructs. Bottom-Right: Vesicles from HEK293T cells transfected with NYESO1/D9/EABR and CD80/EABR constructs. (H) NanoFCM size analysis of vesicles purified by ultracentrifugation from HEK293T cells co-transfected with NYESO1/D9/EABR and CD80/EABR constructs. (I) Immuno-EM image of vesicles purified by ultracentrifugation and size-exclusion chromatography from Expi293F cells co-transfected with NYESO1/D9/EABR and CD80/EABR constructs. Primary human anti-NYESO1:HLA-A*02:01 (3M4E5) antibody and mouse anti-CD80. Secondary 12 nm gold-conjugated anti-mouse antibody appears as black punctae in image and highlighted by green arrow. Secondary 6nm gold-conjugated anti-human antibody appears as smaller black punctae in image and highlighted by blue arrow. Scale bar, 200 nm.

To quantify the relative concentration of the transfected proteins on our engineered vesicles, we performed sandwich ELISAs of the purified APVs using capture antibodies for either the co-expressed immunoregulatory proteins (Figure 2B) or the NYESO1:HLA-A*0201 complex (Figure 2C) followed by biotinylated detection antibodies for the opposite co-presenting protein. For all conditions tested, the best vesicle production occurs when at least one of the transfected proteins contains an EABR sequence (Figure 2D-F). Interestingly, our ELISA data revealed that the ratio of proteins on the surface of the vesicles is influenced not only by which of the two transfected proteins is appended by an EABR sequence, but also by the identity of the co-expressed immunoregulatory protein. To better understand the relative ratio of proteins on a per-vesicle basis, we performed nano flow cytometry (nanoFCM)^40^ of the vesicles from Figure 2F that co-present CD80 and NYESO/HLA-A*0201 (Figure 2G). Using the same antibodies as those used for the ELISA from Figure 2F, the nanoFCM detected a more concentrated presence of CD80 on vesicles when CD80 was appended by an EABR sequence, which would be expected given that both CD80 and pMHCI would then be capable of inducing vesicle formation. Understandably, not all proteins will express the same from DNA transfection; thermostability and sequence-length can influence the ultimate membrane concentration of the protein, so it is not unexpected that addition of an EABR sequence to both pMHCI and CD80 led to an increase in the number of “unlabeled” particles present in quadrant P3. The “unlabeled” P3 quadrant also shows that not all vesicles generated by cell cultures transfected with EABR-tagged proteins feature EABR-tagged proteins on their surface, which would imply that the transfected CD80 and pMHCI proteins are competing with other vesicle-forming proteins or existing, unexploited vesicle-forming pathways for presentation on extracellular vesicles. Measurements of vesicle diameter provided by the nanoFCM demonstrate that most of the particles are between 60-80 nm in size, corroborating our previous APV size measurements^31^ (Figure 2H). Notably, some vesicles are over 100 nm in diameter, which we further confirmed with immuno-electron microscopy (Figure 2I) by co-staining with the same anti-CD80 and anti-NYESO1:HLA-A*0201 antibodies used in Figure 2F and Figure 2G.

Altogether, these results demonstrate that the final protein composition of the vesicles generated using the EABR sequence can be finely tuned depending on the selective addition or removal of EABR from transfected proteins, and that inclusion of the EABR sequence allows for the high-yield synthesis of co-presenting immunoregulatory extracellular vesicles that mimic the endogenous immunoregulatory APVs studied extensively in literature^41, 42^.

### Functionalizing nanovials with vesicles co-presenting pMHCI/EABR and CD80 improves T cell loading and activation

Koo and Mao et al. previously demonstrated that nanovials coated with peptide-major histocompatibility complex (pMHC) monomers could be used to isolate and selectively activate antigen-reactive T cells^12^. While this capture and activation method was successful in isolating previously unidentified reactive T cells, we hypothesized that functionalizing the nanovial surface with EABR-mediated APVs would provide a more physiologically representative interface for the “immunological synapse” that forms between an APC and a T cell during antigen presentation and subsequent immune activation^11^. As a known positive control for comparison, we recreated the original experimental conditions for pMHCI nanovials by coating the interior cavity of nanovials with 20ug/mL biotinylated anti-human IFN-γ and 20ug/mL biotinylated NYESO1-MHCI monomer before incubating with human peripheral blood mononuclear cells (PBMCs) that had been transduced with the 1G4 TCR targeting the NYESO:MHCI complex^43, 44^. Truncated nerve growth factor receptor (NGFR) was used as the cotransduction marker for the presence of the 1G4 TCR, and Calcein AM stain confirmed the cells captured in the nanovials were alive and capable of responding to pMHCI activation with IFN-γ secretion (Figure 3A).

**Figure 3.**
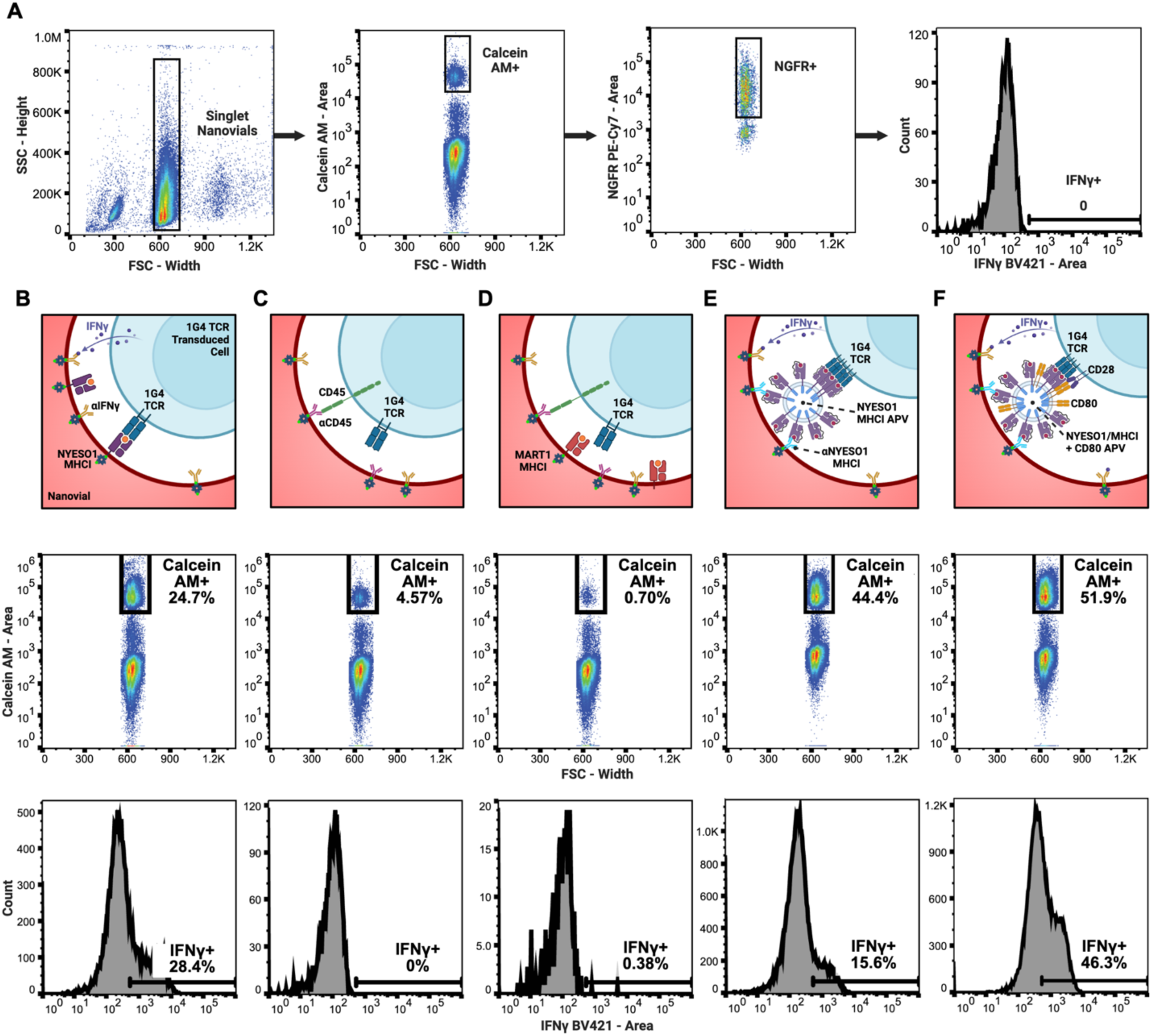
Functionalizing nanovials with vesicles co-presenting pMHCI/EABR and CD80 improves T cell loading and activation. (A) Flow cytometry gating strategy for T-cell loaded nanovials and secretion detection. Singlet nanovials are identified as separate from hydrogel debris and nanovial doublets. Positive Calcein AM signal represents living cells. Positive NGFR signal represents successful transduction of 1G4 TCR. IFN-γ release is measured by an ɑIFN-γ BV421 sandwich antibody. (B-F) First row: Comparison of different pMHC vesicles conjugated to nanovial cavity surface. Second row: Relative cell loading rates into nanovials using different binding moieties. Third row: IFN-γ secretion levels from single T cells on nanovials. T cell loading is significantly improved by using a matched pMHC relative to standard anti-CD45 antibody-based binding or inappropriately matched MART1/HLA-A*0201 for 1G4 TCRs. EABR-mediated pMHC vesicles further improve T cell loading rates. NYESO1/HLA-A*0201 pMHCs in both protein monomer and vesicle format trigger 1G4 T cell activation. (B) Top: Schematic showing nanovials incubated with 20 μg/mL biotinylated NYESO1:HLA-A*0201 monomer and 20 μg/mL biotinylated ɑIFN-γ for 1G4 T cell and IFN-γ capture respectively. Middle: 24.7% of nanovials captured live T cells. Bottom: 28.4% of the live cells captured in nanovials stained positively for IFN-γ release. (C) Top: Schematic showing nanovials incubated with 20 μg/mL biotinylated ɑCD45 and 20 μg/mL biotinylated ɑIFN-γ for 1G4 T cell and IFN-γ capture respectively. Middle: 4.57% of nanovials captured live T cells. Bottom: 0% of the live cells captured in nanovials stained positively for IFN-γ release. (D) Top: Schematic showing nanovials incubated with 20 µg/mL biotinylated MART1:HLA-A*0201 monomer and 20 μg/mL biotinylated ɑIFN-γ for 1G4 T cell and IFN-γ capture respectively. Middle: 0.70% of nanovials captured live T cells. Bottom: 0.38% of the live cells captured in nanovials stained positively for IFN-γ release. (E) Top: Schematic showing nanovials incubated with 20 µg/mL biotinylated ɑNYESO1:HLA-A*0201 antibody and 20 μg/mL biotinylated ɑIFN-γ, followed by incubation with NYESO1/HLA-A*0201/EABR APVs for 1G4 T cell and IFN-γ capture. Middle: 44.4% of nanovials captured live T cells. Bottom: 15.6% of the live cells captured in nanovials stained positively for IFN-γ release. (F) Top: Schematic showing nanovials incubated with 20 µg/mL biotinylated ɑNYESO1:HLA-A*0201 antibody and 20 μg/mL biotinylated ɑIFN-γ, followed by incubation with NYESO1/HLA-A*0201/EABR+CD80/EABR APVs for 1G4 T cell and IFN-γ capture. Middle: 51.9% of nanovials captured live T cells. Bottom: 46.3% of the live cells captured in nanovials stained positively for IFN-γ release.

In accordance with the original flow cytometry experiment by Koo and Mao et al.^12^, we achieved excellent cell capture with live 1G4 T cells occupying ∼25% of the nanovials, of which 28.4% were activated (Figure 3B). By comparison, nanovials coated with 20 μg/mL biotinylated anti-CD45 (Figure 3C), which should capture T cells independent of TCR, or a combination of 20 µg/mL biotinylated anti-CD45 and 20 µg/mL biotinylated MART1-MHCI monomer (Figure 3D), which is a non-cognate pMHCI for 1G4 T cells, showed poor cell capture (4.57% and 0.70% respectively) and activation (0% and 0.38% respectively).

We then captured our EABR-mediated APVs in nanovials by coating the surface with 20 µg/mL biotinylated anti-human NYESO1/HLA-A2 (3M4E5) before incubating with our engineered vesicles (Figure 3E). We intentionally chose the TCR-like anti-human NYESO1/HLA-A2 (3M4E5) antibody, which binds to the TCR binding site on pMHCI^45, 46^, for vesicle capture to confirm that the activation of T cells would be due to intact APVs and not soluble pMHCI monomer from lysed vesicles. Nanovials functionalized with NYESO1/HLA-A2/EABR APVs improved T cell loading to ∼45%, and 15.6% of these cells were activated to release IFN-γ. We used the same 20 µg/mL biotinylated anti-human NYESO1/HLA-A2 (3M4E5) antibody to capture APVs co-presenting NYESO1/HLA-A*0201/EABR and CD80 (Figure 3F); T cell capture increased further to ∼52%, of which an increased fraction of 46.3% were activated to release IFN-γ, a result that would be expected from costimulation by CD80.^47^ This 100% improvement in cell loading is in spite of our use of the single-chain heterotrimer format of pMHCI^37–39^, which should theoretically bind with less affinity than the recombinant, refolded soluble pMHCI used by Koo and Mao et al.; the flexible glycine-serine linker connecting the presenting NYESO1 peptide to β2m travels outward from the closed groove of the MHCI peptide binding cleft toward the TCR, and is thus expected to interfere with optimal TCR binding^38, 39^.

As a result, we conclude that EABR-mediated APVs improve the interface of hydrogel surfaces for T cell capture and activation compared to recombinant protein alone. This improvement in cell binding and activation may enable greater sensitivity and improve the recovery rate of rare cancer-specific functional TCRs for adoptive T cell therapies. Further, we demonstrated that a single antibody can be used to capture a variety of co-presenting immunoregulatory vesicles, which will enable future high-throughput single cell studies of extracellular vesicle protein composition on T cell capture, activation, and differentiation.

### Co-expression of HER2/EABR and CD54 generates vesicles mimicking cancer exosomes

Following the successful functionalization of nanovials with EABR-mediated APVs for the capture of T cells, we hypothesized that we could also capture CAR-T cells by functionalizing the nanovial surfaces with EABR vesicles presenting membrane-bound cancer antigens. Membrane proteins in general have long presented a challenge to biochemical and functional studies as membrane proteins may be insoluble or display altered or absent activities in the absence of a lipid bilayer^17, 18^. For instance, while antibodies screened against recombinant HER2 have been demonstrated to similarly bind membrane-bound HER2, HER2 is known to homodimerize when the protein is overexpressed and densely populated in the lipid bilayer of the cancer cell membrane^48^; in this context, rare HER2 CAR-T cells that specifically bind the homodimer complex may activate more selectively against cancer cells, but these reactive CAR-T cells might otherwise be lost when screening with recombinant HER2. To generate vesicles presenting more natural, membrane-bound HER2, we appended the EABR vesicle-forming sequence to the cytoplasmic tail of the full-length HER2 protein and purified the vesicles from transfected Expi293F cell culture supernatant (Figure 4A). Recent observations on the immunological synapse formed by CAR-T cells has highlighted the significance of membrane-bound CD54 for organizing and maintaining the supramolecular activation complex that leads to T cell activation^49^ or suppression^50^, so we additionally generated vesicles co-presenting both HER2/EABR and CD54 by co-transfecting cell cultures with DNA plasmids encoding both constructs (Figure 4B). An added benefit of co-expressing CD54 was that future cancer-relevant membrane proteins, such as mesothelin^51^, carcinoembryonic antigen^52^, or prostate-specific membrane antigen^53^, could be presented similar to HER2 on the EABR-mediated vesicles and the vesicles could be captured within nanovials by using the same CD54 antibody without requiring a bespoke antibody for every tested cancer antigen.

**Figure 4.**
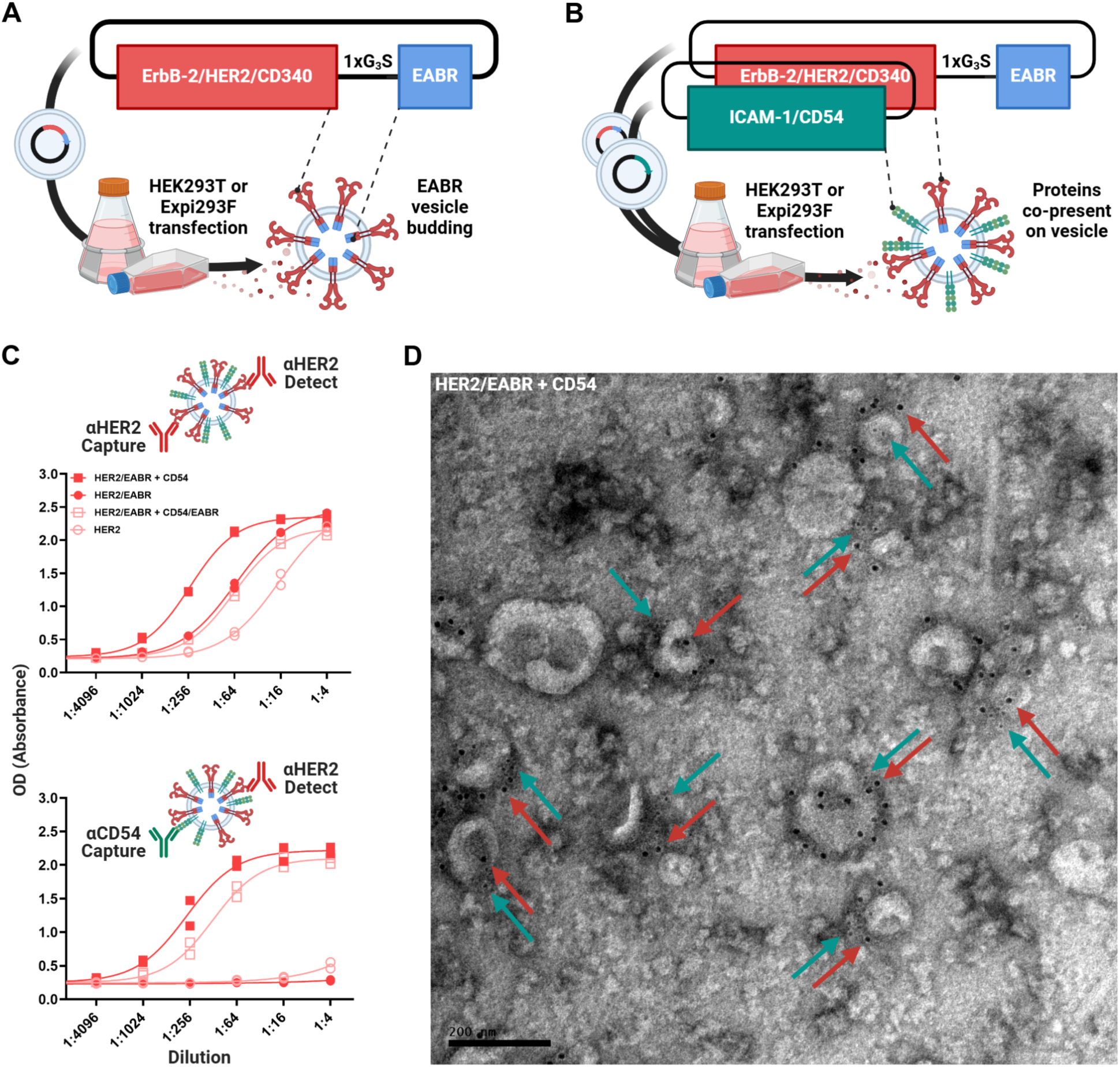
Co-expression of HER2/EABR and CD54 presents both proteins on purified extracellular vesicles. (A) Schematic showing the plasmid construct for HER2 vesicles featuring a G3S spacer and the “EABR” vesicle-forming sequence. The plasmid is transfected into HEK293T or Expi293F cells and vesicles are harvested as shown previously in Figure 1B. (B) Schematic showing the plasmid construct for HER2 + CD54 vesicles. Only HER2 requires a vesicle-forming “EABR” sequence to be appended for both CD54 and HER2 to appear on the harvested vesicles. (C) Sandwich ELISA of APVs purified by ultracentrifugation from transfected Expi293F cells. Cells were co-transfected with HER2/EABR and CD54, HER2/EABR alone, HER2/EABR and CD54/EABR, or HER2 constructs (2 replicates per dilution). Top: anti-HER2 for capture, biotinylated anti-HER2 for detection, and HRP-conjugated streptavidin secondary. Bottom: anti-CD54 for capture, biotinylated anti-HER2 for detection, and HRP-conjugated streptavidin secondary. (D) Immuno-EM image of vesicles purified by ultracentrifugation and size-exclusion chromatography from Expi293F cells co-transfected with HER2/EABR and CD54 constructs. Primary mouse anti-HER2 antibody and rabbit anti-CD54. Secondary 12 nm gold-conjugated anti-mouse antibody appears as black punctae in image and highlighted by red arrows. Secondary 6nm gold-conjugated anti-rabbit antibody appears as smaller black punctae in image and highlighted by blue arrows. Scale bar, 200 nm.

To quantify the relative yield of HER2-presenting vesicles by addition of the EABR sequence, we performed sandwich ELISAs with either HER2 antibodies or a sandwich of CD54 capture antibody and HER2 detection antibody (Figure 4C). In agreement with previous studies on HER2-presenting exosomes that originate from HER2 over-expressing cancer cells^54^, over-expression of the full-length HER2 sequence in high-expressing Expi293F cells spontaneously generated vesicles without the addition of an appended EABR sequence. However, addition of the EABR sequence to HER2 further improved the yield of HER2+ vesicles. Interestingly, CD54 seems to act synergistically with HER2 to boost HER2 vesicle production, as co-transfection with separate DNA plasmids encoding HER2/EABR (5ug) and CD54 (5ug) leads to better vesicle expression than transfection with 10ug of plasmid DNA encoding HER2/EABR alone. Appending EABR to both HER2 and CD54 leads to a reduction in total HER2 detected on the purified vesicles, which is likely due to an increase in CD54 presentation on the vesicles at the expense of HER2 presentation, a phenomenon that was observed previously when co-transfecting DNA constructs encoding equal amounts of NYESO1/HLA-A*0201I/EABR and CD80/EABR (Figure 2G). We further confirmed that the detected co-presentation of HER2 and CD54 is localized on vesicle membranes by performing immuno-EM of the supernatant from transfected cells (Figure 4D). Simultaneous co-staining of αCD54 and αHER2 showed halos of black punctae surrounding exosome-like vesicles mimicking cancer exosomes. Because our goal was to maximize HER2 presentation on vesicles while co-presenting a generalizable “capture” protein on the vesicle membrane, we moved forward with subsequent nanovial characterization experiments using the co-presenting HER2/EABR+CD54 vesicles as well as the HER2/EABR vesicles for comparison.

### Functionalizing nanovials with vesicles co-presenting HER2/EABR and CD54 improves CAR-T cell loading and activation

To optimize the capture of HER2/EABR vesicles inside the nanovial cavity, we incubated the nanovials with different concentrations of αHER2 capture antibody to determine the minimal concentration necessary for maximal HER2 signal as detected by a fluorophore-conjugated αHER2 sandwich antibody. HER2 signal from captured HER2/EABR vesicles was saturated at 20 ug/mL αHER2 capture antibody (Figure 5A), a concentration similar to that of the pMHCI nanovials. Surprisingly, this maximum fluorescent signal from HER2/EABR vesicles was orders of magnitude lower than the nanovials functionalized with biotinylated recombinant HER2. In comparison, coating the nanovials with biotinylated αCD54 capture antibodies and subsequent incubation with co-presenting HER2/EABR+CD54 vesicles produced an αHER2 fluorescent signal that was equivalent to biotinylated recombinant HER2 (Figure 5B). As was demonstrated by previous nanoFCM analysis, immuno-EM micrographs, and ELISA results of EABR vesicles, we believe this dramatic increase in signal between the HER2/EABR and co-presenting HER2/EABR+CD54 vesicles is likely due to differences in the binding affinities of the αHER2 and αCD54 capture antibodies for their target proteins, and not due to differences in vesicle stability or the relative density of HER2 or CD54 protein on the vesicle surface. Nonetheless, the comparable level of HER2 signal that was detected between the recombinant HER2 and HER2/EABR+CD54 vesicles captured within the nanovials provided a convenient comparison for our ensuing studies on the relative importance of a lipid bilayer and CD54 co-presentation for T cell capture and activation.

**Figure 5.**
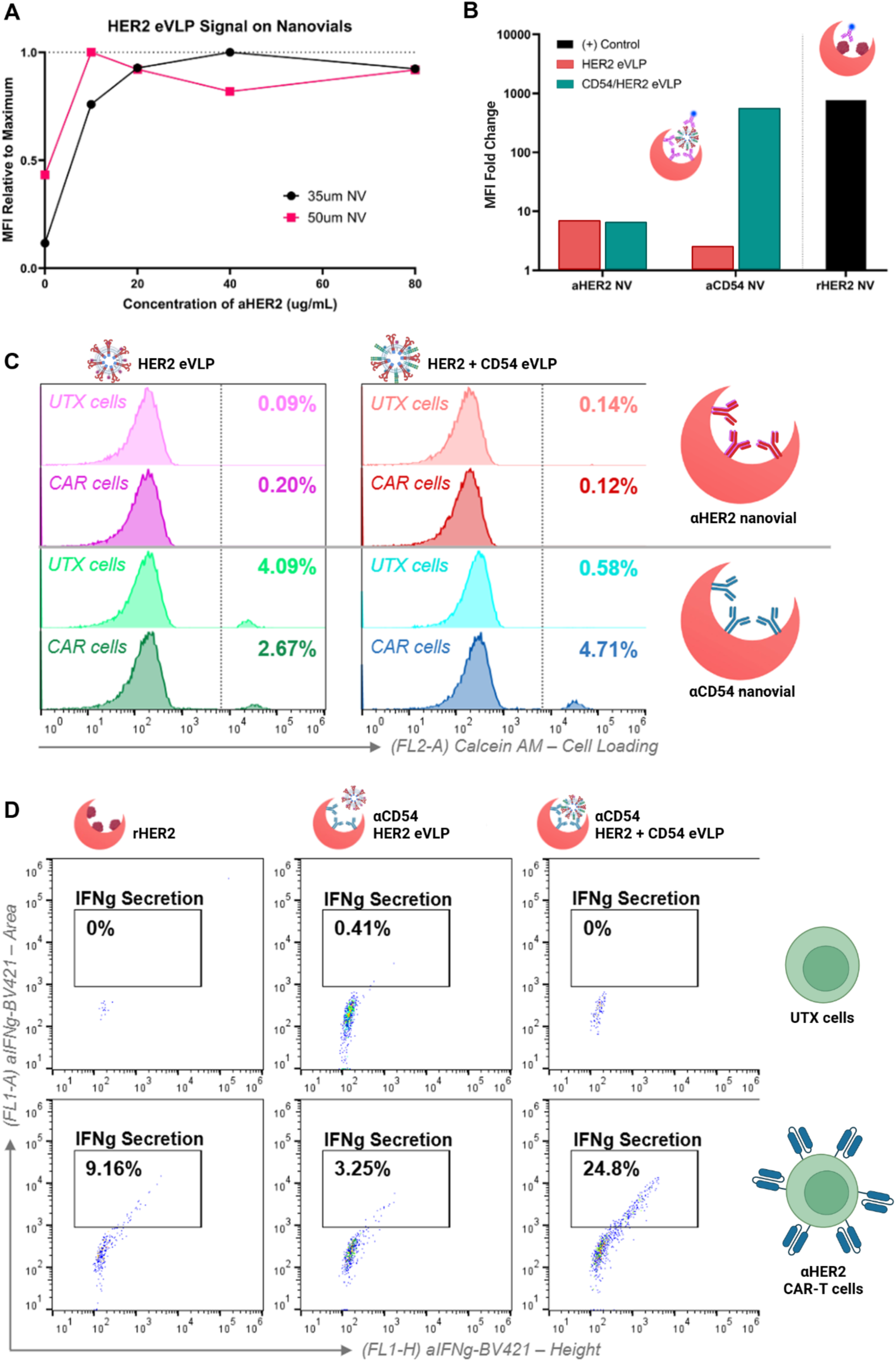
Functionalizing nanovials with vesicles co-presenting HER2/EABR and CD54 improves CAR-T cell loading and activation. (A) Immobilization of HER2 vesicles on 35um and 50um nanovials based on concentration of ɑHER2 capture antibodies on the nanovials. Maximum capture signal is detected at 20 ug/mL ɑHER2 for both 35um and 50um nanovials. (B) HER2/EABR+CD54 vesicles show significantly higher capture rate on nanovials compared to the HER2-only vesicles. This signal is comparable to the positive control of nanovials conjugated with 20 ug/mL recombinant HER2 (rHER2). (C) ɑHER2 CAR T cell and control untransfected (UTX) T cell loading in nanovials using both HER2/EABR vesicles and HER2/EABR+CD54 vesicles. Nanovials conjugated with ɑHER2 antibodies, then incubated with either HER2/EABR or HER2/EABR+CD54 vesicles, cannot bind control UTX cells nor ɑHER2 CAR T cells. Nanovials conjugated with ɑCD54 antibodies show non-specific cell loading which may be due to endogenous CD54 expression on T cells. This non-specific binding is significantly reduced when HER2/EABR+CD54 vesicles are immobilized on nanovials. Nanovials conjugated with ɑCD54 antibodies, then incubated with HER2/EABR+CD54 vesicles, bind ɑHER2 CAR T cells selectively. (D) Flow data showing improvement in nanovial assay from using EABR-mediated vesicles. Nanovials functionalized with rHER2 show 9.16% of captured, living CAR T cells are activated to release IFN-γ. Nanovials functionalized with ɑCD54 antibodies, then incubated with HER2/EABR+CD54 vesicles, show 24.8% of captured, living CAR T cells are activated to release IFN-γ.

To investigate the significance of a lipid bilayer and co-presentation of CD54 on T cell capture and activation by cancer antigens, we compared the efficiency of nanovial loading between untransfected (UTX) T cells and T cells transfected with an αHER2 CAR (Figure 5C). Nanovials that were conjugated with biotinylated αHER2 antibodies and incubated with either HER2/EABR or HER2/EABR+CD54 vesicles did not appreciably bind UTX cells nor αHER2 CAR T cells, which aligned with our previous tests of HER2 signal intensity in nanovials (Figure 5B) and would support our conjecture that our chosen αHER2 antibody has low affinity for HER2. Alternatively, nanovials that were conjugated with biotinylated αCD54 antibodies and incubated with HER2/EABR+CD54 vesicles bound αHER2 CAR T cells selectively; these functionalized nanovials also showed a dramatic increase in CAR T cell activation compared to recombinant HER2 as measured by the levels of IFN-γ release and fraction of cells releasing IFN-γ (Figure 5D). Nanovials conjugated with biotinylated αCD54 antibodies and incubated with HER2/EABR vesicles bound both αHER2 CAR T cells and UTX T cells indiscriminately; this non-specific binding of UTX cells is significantly reduced when HER2/EABR+CD54 vesicles are instead immobilized on the αCD54 nanovials, suggesting that the HER2/EABR+CD54 vesicles are saturating the αCD54 antibodies present on the nanovial surface and preventing binding to possible endogenous CD54 expression on T cells.

Our results demonstrate that EABR-mediated cancer-mimicking vesicles that co-present CD54 and HER2 improve CAR T cell capture and activation on hydrogel surfaces. Antibody affinity is a limiting factor for vesicle functionalization on hydrogel surfaces, but, as we demonstrated, CD54 co-expression enables convenient and generalizable functionalization of hydrogel surfaces with EABR-mediated vesicles that present the natural structure of cancer antigens for high-throughput capture, activation, and downstream analysis of therapeutically-relevant CAR T cells.

## Discussion

Here, we present a method for improving the T cell activation potential and T cell capture rates of the artificial antigen-presenting cavities in nanovials by functionalizing their hydrogel surface with engineered EABR-mediated vesicles mimicking APVs and cancer exosomes. EABR-mediated vesicles are densely coated in transfected immunoregulatory proteins embedded in a natural, cell-derived lipid bilayer, which provides a more physiologically representative interface for the “immunological synapse” that forms between an APC and a T cell during antigen presentation and subsequent immune activation. This approach can increase the sensitivity and control of nanovial-based enrichment of natural and artificial libraries of T cells and CAR-T cells to identity and isolate reactive antigen-specific T cells on a single-cell basis. Microparticle hydrogel nanovials functionalized with the EABR-mediated vesicles show a two-fold improvement in T cell capture of 1G4 T cells and an over two-fold improvement in activation of HER2 CAR-T cells compared to nanovials functionalized with recombinantly-expressed soluble protein. Further, we demonstrated the ability to fine-tune the composition of multiple co-presenting immunoregulatory proteins on single particles to more accurately reflect, or systematically modulate, the components of the cell surface of an APC. Future studies can either repurpose our methods for capturing endogenous EVs on nanovials or synthesize EABR-mediated immunoregulatory extracellular vesicles to study the influence of EV protein composition on T cell activation and differentiation.

As observed by the synergistic increase in vesicle release when both HER2/EABR and CD54 constructs were co-transfected, a future point of optimization for EV synthesis is seeing what parameters can be further engineered to maximize vesicle production. The vast majority of vesicles we detected by nanoFCM did not feature an EABR-tagged protein, implying that transfected EABR-proteins compete with endogenous vesicle-forming proteins or existing, unexploited vesicle-forming pathways for presentation on extracellular vesicles. Similarly, the range of vesicles detected by nano-FCM suggests that it may be possible to control the size distribution of the vesicles. Previous studies have demonstrated an improvement in T cell activation when synthetic antigen-presenting nanoparticles are larger in diameter^55^, but it is unclear how vesicle size will influence T cell activation when the vesicles are immobilized as a collection on the surface of a hydrogel. The improvement in T cell capture and activation we observed with our EABR-mediated vesicles compared to recombinant protein may be due to the density of stimulatory cues on the EABR-mediated vesicles^29^, the fluid lipid bilayer^11, 14^, the co-presentation of cell-membrane derived proteins like CD54^49, 50^, or the presence of the natural full-length membrane proteins^18^. Future studies that isolate the critical elements of T cell activation by these EABR-mediated vesicles will lead to more optimal hydrogel functionalization strategies for T cell activation. Nanovial-based approaches may also be useful to select cells that secrete more extracellular vesicles, as was previously done for natural extracellular vesicles released by mesenchymal stem cells.

In summary, our current work here expands upon the growing collection of characterization studies and workflows already pioneered by nanovials^12, 13, 56–59^, which allow standard flow sorters and single-cell sequencing instrumentation to accelerate the development of personalized cell therapies. Additionally, with respect to the growing interest in cell-free, cell-like therapies^60–63^, we demonstrate here a system for optimizing the engineering and characterization of extracellular vesicles that can be used either cooperatively^64^ or in place of adoptive cell therapies^65, 66^ for targeted immunoregulation.

## Acknowledgements

We would like to thank other members of Di Carlo lab and Witte lab for helpful comments and discussion in preparation of this manuscript. We thank UCLA Jonsson Comprehensive Cancer Center (JCCC) and Center for AIDS Research Flow Cytometry Core Facility. We thank J. Vielmetter, A. Lam, and the Caltech Protein Expression Center for assistance with protein production; P. Gnanapragasam, A. Rorick, L. Segovia for cell culture reagents; W. Chour for MHC plasmid maps and helpful discussions. We thank R. Voorhees and J. Keeffe for equipment access and training. Immuno-electron microscopy was performed in the Caltech Cryo-EM Center with assistance from Songye Chen. Figures 1A-D, 2A-C, 2G, 3B-F, 4A-C, and 5B-D were created with Biorender.com. Research was sponsored by the U.S. Army Research Office and accomplished under cooperative agreement W911NF-19-2-0026 for the Institute for Collaborative Biotechnologies. The content of the information on this page does not necessarily reflect the position or policy of the Government, and no official endorsement should be inferred. This project has been made possible in part by grant 2023-332386 from the Chan Zuckerberg Initiative Donor Advised Fund (CZI DAF), an advised fund of the Silicon Valley Community Foundation.

## Author Contributions

B.A.O., M.M., C.S., Z.M., D.D.C., and S.L.M. designed research; B.A.O., M.M., C.S., and K.L. performed research; A.M. and Z.M. contributed new reagents/analytic tools; B.A.O., M.M., C.S., Z.M., K.L., M.A.G.H., D.D.C., and S.L.M. analyzed data; B.A.O. wrote the manuscript; all authors edited the manuscript.

## Corresponding authors

Further information and requests for resources and reagents should be directed to and will be fulfilled by the lead contact, Stephen L. Mayo.

## Materials availability

All expression plasmids generated in this study are available upon request through a Materials Transfer Agreement.

## Data and code availability

All data are available in the main text. Materials are available upon request to the corresponding authors with a signed material transfer agreement. Any additional information required to reanalyze the data reported in this paper is available from the lead contact upon request. This paper does not report original code. This work is licensed under a Creative Commons Attribution 4.0 International (CC BY 4.0) license, which permits unrestricted use, distribution, and reproduction in any medium, provided the original work is properly cited. To view a copy of this license, visit https://creativecommons.org/licenses/by/4.0/. This license does not apply to figures/photos/artwork or other content included in the article that is credited to a third party; obtain authorization from the rights holder before using such material.

## Methods

### Key resources table

**Table S1.** CryoEM data acquisition and image processing.

| REAGENT or RESOURCE | SOURCE | IDENTIFIER |
| --- | --- | --- |
| **Antibodies** |  |  |
| Recombinant Anti-HLA-A2-peptide (SLLMWITQV) Complex (3M4E5) Mouse Monoclonal Antibody | ProteoGenix | Reference:<br>RHM03002-100 |
| Purified anti-human CD340 (erbB2/HER-2) Antibody | Biolegend | Cat# 324402<br>RRID: AB_756118 |
| ICAM-1 Recombinant Rabbit Monoclonal Antibody (ST0487) | Thermo Fisher Scientific | Cat# MA5-41137<br>RRID: AB_2898891 |
| CD54 (ICAM-1) Monoclonal Antibody (HA58) | Thermo Fisher Scientific | Cat# 14-0549-82<br>RRID: AB_467304 |
| ErbB2 (HER-2) Monoclonal Antibody (3B5), Biotin | Thermo Fisher Scientific | Cat# MA5-13672<br>RRID: AB_10982527 |
| CD54 (ICAM-1) Monoclonal Antibody (HA58), Biotin | Thermo Fisher Scientific | Cat# 13-0549-82<br>RRID: AB_466483 |
| CD80 (B7-1) Monoclonal Antibody (2D10.4) | Thermo Fisher Scientific | Cat# 14-0809-82<br>RRID: AB_467350 |
| CD80 (B7-1) Monoclonal Antibody (2D10.4), Biotin | Thermo Fisher Scientific | Cat# 13-0809-82<br>RRID: AB_466513 |
| Purified anti-human CD83 Antibody | Biolegend | Cat# 305302<br>RRID: AB_314509 |
| Biotin anti-human CD83 Antibody | Biolegend | Cat# 305304<br>RRID: AB_314511 |
| CD274 (PD-L1, B7-H1) Monoclonal Antibody (MIH1) | Thermo Fisher Scientific | Cat# 14-5983-82<br>RRID: AB_467784 |
| CD274 (PD-L1, B7-H1) Monoclonal Antibody (MIH1), Biotin | Thermo Fisher Scientific | Cat# 13-5983-82<br>RRID: AB_466840 |
| 6 nm Colloidal Gold AffiniPure® Goat Anti-Human IgG (H+L) (EM Grade) | Jackson ImmunoResearch | Cat# 109-195-088<br>RRID: AB_2337747 |
| 6 nm Colloidal Gold AffiniPure® Goat Anti-Rabbit IgG (H+L) (EM Grade) | Jackson ImmunoResearch | Cat# 111-195-144<br>RRID: AB_2338015 |
| 12 nm Colloidal Gold AffiniPure® Goat Anti-Mouse IgG (H+L) (EM Grade) | Jackson ImmunoResearch | Cat# 115-205-146<br>RRID: AB_2338733 |
| anti-IFN-γ ms mAb – BV421 | BioLegend | Cat# 502532 |
| anti-HER2/ErbB2 Antibody – biotin | Sino Biological | Cat# 10004-MM01-B |
| Biotin anti-human CD45 Ab, 2D1 | BioLegend | Cat# 368534 |
| Human IFN-γ Biotinylated Antibody | Biotechne/R&D Systems | Cat# BAF285 |
| Anti-CD340 (erbB2/HER2) ms mAb – PE/Cy7 | BioLegend | Cat# 324413 |
| **Bacterial and virus strains** |  |  |
| *E. coli* DH5 Alpha | Zymo Research | Cat# T3009 |
| **Chemicals, peptides, and recombinant proteins** |  |  |
| Luria Broth Base (Miller's LB Broth Base), powder | Thermo Fisher Scientific | Cat# 12795084 |
| Dulbecco's Modified Eagle Medium (DMEM) | Gibco | Cat# 11995-065 |
| Fetal bovine serum (FBS) | R&D Systems | Cat# S12450H |
| Penicillin-Streptomycin | Gibco | Cat# 15-140-122 |
| FuGENE® HD Transfection Reagent | Promega | Cat# E2311 |
| OPTI-MEM I Reduced Serum Medium | Gibco | Cat# 31985-062 |
| Expi293 Expression Medium | Gibco | Cat# A14351-01 |
| Expi293 Transfection Kit | Gibco | Cat# A14524 |
| NucleoBond™ Xtra Maxi Plus | Macherey-Nagel | Cat# 12758412 |
| Zyppy Plasmid Miniprep Kit | Zymo Research | Cat# D4019 |
| Sucrose | Sigma-Aldrich | Cat# S5016-1KG |
| NaCl | Fisher Scientific | Cat# BP358-212 |
| Sodium hydroxide | Macron Chemicals | Cat# 7708-10 |
| Sodium bicarbonate | Macron Chemicals | Cat# 7412-12 |
| Sodium phosphate dibasic | Sigma-Aldrich | Cat# 04272-1KG |
| Tris-HCl | Fisher Scientific | Cat# BP153-1 |
| TWEEN® 20 | Sigma-Aldrich | Cat# P1379-500ML |
| BSA | Sigma-Aldrich | Cat# B4287-5G |
| Trypsin | Gibco | Cat# 25200-056 |
| L-glutamine | Gibco | Cat# 25030-149 |
| Phytohemagglutinin, M form | Gibco | Cat# 10576015 |
| Paraformaldehyde 16% Aqueous Solution EM Grade | Electron Microscopy Sciences | Cat# 15710 |
| Uranyl Formate | Electron Microscopy Sciences | Cat# 22451 |
| 1-Step™ Ultra TMB-ELISA Substrate Solution | Thermo Fisher Scientific | Cat# 34029 |
| EZ-Link NHS-PEG4 Biotinylation Kit | Thermo Fisher Scientific | Cat# 21455 |
| Streptavidin (HRP) | Abcam | Cat# AB7403 |
| Sulfo-NHS-Biotin | ApexBio | Cat# A8001 |
| Streptavidin | Thermo Fisher Scientific | Cat# 434302 |
| Human HER2/ErbB2 Protein (ECD), Biotinylated | Sino Biological | Cat# 10004-HCCH-B |
| X-VIVO15 Serum-free Media | Lonza | Cat# 04-418Q |
| Invitrogen Molecular Probes Calcein, AM | Thermo Fisher Scientific | Cat# C3099 |
| **Critical commercial assays** |  |  |
| **Experimental models: Cell lines** |  |  |
| HEK293T cells | 67 | RRID:CVCL_0063 |
| Expi293F cells | Gibco | RRID:CVCL_D615 |
| **Recombinant DNA** |  |  |
| HLA-A2-peptide (SLLMWITQV) Complex (3M4E5) Human Monoclonal Antibody heavy chain | 68 | https://patentscope.wipo.int/search/en/detail.jsf?docId=EP32586798 |
| HLA-A2-peptide (SLLMWITQV) Complex (3M4E5) Human Monoclonal Antibody light chain | 68 | https://patentscope.wipo.int/search/en/detail.jsf?docId=EP32586798 |
| D9 SCT MART1/ HLA-A*0201/EABR | This paper | N/A |
| D9 SCT NYESO1/ HLA-A*0201 | This paper | N/A |
| D9 SCT NYESO1/ HLA-A*0201/EABR | This paper | N/A |
| CD80 | This paper | N/A |
| CD80/EABR | This paper | N/A |
| CD83 | This paper | N/A |
| CD83/EABR | This paper | N/A |
| CD274 | This paper | N/A |
| CD274/EABR | This paper | N/A |
| HER2 | This paper | N/A |
| HER2/EABR | This paper | N/A |
| CD54 | This paper | N/A |
| CD54/EABR | This paper | N/A |
| **Software and algorithms** |  |  |
| Image J 1.54f | NIH | https://imagej.net/<br>RRID: SCR_003070 |
| Prism 10.3.1 | GraphPad |  |
| **Other** |  |  |
| Amicon Ultra-15 100 kDa MWCO Centrifugal Filter Unit | EMD Millipore | Cat# UFC910096 |
| Amicon Ultra-4 10 kDa MWCO Centrifugal Filter Unit | EMD Millipore | Cat# UFC801024 |
| Nanosep with 100K Omega | Pall Life Sciences | Cat# OD100C34 |
| Erlenmeyer cell culture flasks - 500mL | Corning | Cat# 431145 |
| 6-well Clear Flat Bottom TC-treated Multiwell Cell Culture Plate, with Lid | Falcon | Cat# 353046 |
| 30 mL BD Luer-Lok™ Syringe | BD | Cat# 302832 |
| 3 mL BD Luer-Lok™ Syringe | BD | Cat# 309657 |
| AirOtop enhanced seal for 125 ml ultra yield flask | VWR | Cat# 899421 |
| Costar® Spin-X® Centrifuge Tube Filters, 0.22 µm Pore CA Membrane | Corning | Cat# 8160 |
| Polycarbonate centrifuge tubes | Beckman Coulter | Cat# 349622 |
| 0.45µm syringe filter | Corning | Cat# 431220 |
| 0.2µm syringe Filter | VWR | Cat# 28143-300 |
| Nalgene™ Rapid-Flow™ Sterile Disposable Filter Unit - 0.45µm | Thermo Fisher Scientific | Cat# 156-4045 |
| Costar high binding plates, 96-well | Corning | Cat# 9018 |
| TempPlate Sealing Foil | USA Scientific | Cat# 2923-0100 |
| HiTrap MabSelect | Cytiva | Cat# 28408253 |
| Superose 6 Increase 10/300 column | Cytiva | Cat# 29-0915-96 |
| HiLoad 16/600 Superdex 200 column | Cytiva | Cat# 28-9893-35 |
| Formvar/Carbon 300 Mesh grids | Electron Microscopy Sciences | Cat# FCF300-Cu-50 |

### Experimental Model and Subject Details

#### Bacteria

*E. coli* DH5 Alpha cells (Zymo Research) used for expression plasmid productions were cultured in LB broth (Sigma-Aldrich) with shaking at 225 rpm at 37 °C. Plasmids were purified for transfection using Zymogen miniprep kits (Zymo Research) or maxiprep kits (Macherey-Nagel).

#### Cell lines

HEK293T cells plated in 6-well plates (Falcon) were cultured in Dulbecco’s modified Eagle’s medium (DMEM, Gibco) supplemented with 10% heat-inactivated fetal bovine serum (FBS, R&D Systems) and 1 U/mL penicillin-streptomycin (Gibco) at 37 °C and 5% CO_2_. Transfections were carried out with FuGENE transfection reagent (Promega) diluted in Opti-MEM (Gibco).

pi293F cells (Gibco) for APV expression and antibody expression were maintained at 37 °C and 8% CO_2_ in Expi293 expression medium (Gibco). Transfections were carried out with an Expi293 Expression System Kit (Gibco). Falcon tubes sealed with AirOtops (VWR) or culture flasks (Corning) containing Expi293F cells were maintained under shaking at 470 rpm for 10-30 mL transfections and 130 rpm for transfections larger than 30 mL. All cell lines were derived from female donors and were not specially authenticated.

#### T cells

T cells were isolated from human donor whole blood samples by negative selection using the RosetteSep Human T Cell Enrichment Cocktail Kit (STEMCELL Technologies). Isolated T cells were seeded in a fresh complete ImmunoCult™-XF T cell expansion medium (STEMCELL Technologies) at 1 x 10^6^ cells/mL with 2 μL/mL ImmunoCult™ human CD3/CD28 T cell activator. T cells were activated for 3 days and expanded for up to 12 days by changing into fresh expansion medium every 2-3 days. All cells were cultured in incubators at 37 °C and 5% CO_2_.

## Methods

### Design of EABR constructs

The EABR sequence was identical to the sequence used by Hoffmann et al. to create SARS-CoV2-S eVLPs^30^. Briefly, the EABR domain (residues 160-217) of the human CEP55 protein was fused to the C-terminus of full length membrane proteins separated by a (Gly)_3_Ser (GS) linker to generate constructs inserted in a p3BNC expression plasmid. To generate the SCT pMHCI construct, the N-terminus of the HLA-A*0201 alpha chain was extended with a 4x(G_4_S) linker connecting to the C-terminus of B2M, which was itself extended at its N-terminus by a 3x(G_4_S) linker connecting to either the 10-mer MART1 (ELAGIGILTV) or 9-mer NYESO1 (SLLMWITQV) cancer-related peptides to make a single-chain heterotrimer (SCT)^69^. To generate the “D9” version of SCT pMHCI, the alpha heavy chain was further edited with Y84C and A139C substitutions^37–39^. Constructs encoding CD80/EABR, CD83/EABR, CD274/EABR, HER2/EABR, and CD54/EABR all similarly appended an EABR domain onto the full-length native CD80, CD83, CD274, HER2, or CD54 proteins respectively.

### Production of EABR APVs

EABR APVs were generated by one of two means: 1) transfecting Expi293F cells (Gibco) cultured in Expi293F expression media (Gibco) on an orbital shaker at 37 °C and 8% CO2 with plasmid DNA pre-filtered through 0.22 µm Spin-X filters (Corning); 2) transfecting HEK293T cells cultured in Dulbecco’s modified Eagle’s medium (DMEM, Gibco) supplemented with 10% heat-inactivated fetal bovine serum (FBS, Sigma-Aldrich) and 1 U/ml penicillin-streptomycin (Gibco) at 37 °C and 5% CO_2_. 72 hours post-transfection, cells were centrifuged at 1000 x g for 10 min, supernatants were passed through a 0.45 um filter (Corning) with Luer-Lok syringes (BD) or a 0.45 um vacuum filter (Thermo Fisher Scientific), and concentrated using Amicon Ultra-15 centrifugal filters with 100 kDa molecular weight cut-off (Millipore). APVs were purified by ultracentrifugation at 50,000 rpm (135,000 x g) for 2 hours at 4 °C using a TLA100.3 rotor and an OptimaTM TLX ultracentrifuge (Beckman Coulter) on a 20% w/v sucrose cushion in polycarbonate centrifuge tubes (Beckman Coulter). Supernatants and the sucrose cushion were removed, and pellets were re-suspended in 200 uL sterile pH 7.4 PBS at 4 °C overnight. To remove residual cell debris, re-suspended samples were transferred to microcentrifuge tubes and centrifuged at 10,000 x g for 10 min; clarified supernatants were collected for subsequent ELISA and nanoFCM assays. For immuno-EM grid preparations, APVs were further purified by SEC using a Superose 6 Increase 10/300 column (Cytiva) equilibrated with pH 7.4 PBS. Peak fractions in the void volume corresponding to pMHCI APVs were combined and concentrated to 250-500 uL in Amicon Ultra-4 centrifugal filters with 100 kDa molecular weight cut-off (Millipore) or Nanosep with 100K Omega centrifugal filters (Pall). Samples were stored at 4 °C and imaged directly after purification.

### Transduction of TCRs in PBMC

Engineering of candidate TCRs in PBMC was performed according to previous publication.^35^

### Antibody Expression

Anti-HLA-A2-peptide (SLLMWITQV) Complex (3M4E5) antibody was made in-house by expressing the heavy chain and light chain of the 3M4E5 antibody at a 1.5:1 plasmid ratio in Expi293F cells.^33^ Supernatant from the transiently-transfected Expi293F cells (Gibco) was separated from the cells by pelleting at 3500g for 15 minutes and filtering the clarified supernatant through a 0.2 um syringe filter (VWR) or 0.45 um vacuum filter unit (Thermo Fisher Scientific). The filtered supernatant was further purified using Fc-affinity chromatography (HiTrap MabSelect, Cytiva) and SEC (HiLoad 16/600 Superdex 200 column, Cytiva). Peak fractions corresponding to purified antibody proteins were pooled, concentrated, and stored at 4 °C. Biotinylated antibodies for ELISAs were generated and purified using an EZ-Link biotinylation kit (Thermo Fisher Scientific). The in-house generated antibody was compared to a commercially available 3M4E5 antibody (RHM03002-100; ProteoGenix) with ELISA and showed identical performance (data available upon request).

### Immuno-EM of EABR-mediated vesicles

SEC-purified EABR-mediated vesicles were prepared on grids for negative stain transmission electron microscopy at room temperature. Formvar/Carbon 300 Mesh grids (Electron Microscopy Sciences) were first glow discharged. 20 uL of purified sample was pipetted onto paraffin, glow discharged formvar/carbon 300 mesh grids were placed on the droplet of sample for 5 minutes, then gently wicked away by dabbing the edge of the grid on filter paper. The grids were then placed on a 20uL droplet of 1% paraformaldehyde (Electron Microscopy Sciences) in pH 7.4 PBS for 10 minutes, and subsequently wicked away. Grids were washed three times by placing the grids on droplets of pH 7.4 PBS for 5 minutes, and wicking away the PBS with filter paper after each wash. Grids were then blocked by placing the grids on droplets of 5% FBS diluted in pH 7.4 PBS for one hour. Blocking solution was wicked away from the grids, and the grids were then placed on droplets of primary antibody (pMHCI vesicles: αNYESO1:HLA-A*02:01 (3M4E5) diluted to 150ug/mL and mouse αCD80 (#14-0809-82; Thermo Fisher Scientific) diluted 1:10; HER2 vesicles: aCD54 (#MA5-41137; Thermo Fisher Scientific) diluted 1:10 and aHER2 (#324402; BioLegend) diluted 1:20) in 5% sucrose, 5% FBS pH 7.4 PBS for 2 hours. To prevent drying of the primary antibody solution, the grids were placed in a humidified chamber made from wet paper towels folded under a pyrex glass dish. Grids were again washed three times by placing the grids on droplets of pH 7.4 PBS for 10 minutes, and wicking away the PBS with filter paper after each wash. The grids were then placed on droplets of colloidal gold-conjugated secondary antibody (pMHCI vesicles: 6 nm colloidal gold goat anti-human antibody (#109-195-088; Jackson ImmunoResearch) diluted 1:20 and 12 nm colloidal gold goat anti-mouse antibody (#115-205-146; Jackson ImmunoResearch) diluted 1:20; HER2 vesicles: 6 nm colloidal gold goat anti-rabbit antibody (#111-195-144; Jackson ImmunoResearch) diluted 1:20 and 12 nm colloidal gold goat anti-mouse antibody (#115-205-146; Jackson ImmunoResearch) diluted 1:20) in 5% sucrose, 5% FBS pH 7.4 PBS for 1 hour. Grids were subsequently washed three times, ten minutes per wash with pH 7.4 PBS, wicking away the PBS after the ten minute wash each time. Finally, the grid was placed on a droplet of 1.5% uranyl formate (Electron Microscopy Sciences), prepared fresh within 1 week of use, for 1 minute. The excess uranyl formate was wicked away and the grid was left to air dry for 30 minutes before storing at room temperature away from light. Imaging of HER2 grids were performed on a FEI Tecnai T12 Transmission Electron Microscope within two days after the grids were prepared. The NYESO1/HLA-A*0201/EABR+CD80 formvar grids were unexpectedly sensitive to the T12 beam and frequently ruptured before we could acquire an image, so these negative-stain grids were imaged using an FEI Talos Arctica.

### Sandwich ELISA

96-well plates (Corning) were coated with 5 ug/mL of capture antibody diluted in sterile 0.1 M NaHCO_3_ pH 9.6, and sealed with TempPlate sealing foil (USA Scientific) overnight at 4 °C. Capture antibodies included: CD340 (erbB2/HER-2) (#324402; Biolegend), CD54 (ICAM-1) (#14-0549-82; Thermo Fisher Scientific), CD80 (B7-1) (#14-0809-82; Thermo Fisher Scientific), CD83 (#305302; Biolegend), CD274 (PD-L1, B7-H1) (#14-5983-82; Thermo Fisher Scientific). Plates were emptied and blocked with TBS-T/3% BSA for at least 30 minutes. Plates were again emptied, and vesicle samples 4-fold serially diluted in TBS-T/3% BSA were added to the plates. After a 2-hour incubation at room temperature, plates were emptied, and 5 ug/mL of biotinylated detection antibody diluted in TBS-T/3% BSA was added to the plates. Detection antibodies included: biotinylated αErbB2 (HER-2) (#MA5-13672; Thermo Fisher Scientific), biotinylated αCD54 (ICAM-1) (#13-0549-82; Thermo Fisher Scientific), biotinylated αCD80 (B7-1) (#13-0809-82; Thermo Fisher Scientific) biotinylated αCD83 (#305304; Biolegend), biotinylated αCD274 (PD-L1, B7-H1) (#13-5983-82; Thermo Fisher Scientific). After another 2-hour incubation at room temperature, plates were washed three times with TBS-T. HRP-conjugated streptavidin (Abcam) was diluted to manufacturer’s recommendations in TBS-T/3% BSA and added to plates for 30 minutes at room temperature. After washing three times with TBS-T, plates were developed using 1-Step™ Ultra TMB-ELISA Substrate Solution (Thermo Fisher Scientific) and absorbance was measured at 450 nm. Standard 4-parameter sigmoidal binding curves were calculated using GraphPad Prism 10.3.1 without any further editing.

### Nanovial Fabrication

Polyethylene glycol biotinylated nanovials with 35 μm diameters were fabricated using a three-inlet flow-focusing microfluidic droplet generator, sterilized and stored at 4 °C in Washing Buffer consisting of Dulbecco’s Phosphate Buffered Saline (Thermo Fisher) with 0.05% Pluronic F-127 (Sigma), 1% 1X antibiotic-antimycotic (Thermo Fisher), and 0.5% bovine serum albumin (Sigma) as previously reported^13^.

### Nanovial Functionalization

**Streptavidin conjugation to the biotinylated cavity of nanovials.** Sterile nanovials were diluted in Washing Buffer five times the volume of the nanovials (i.e., 100 µL of nanovial volume was resuspended in 400 µL of Washing Buffer). A diluted nanovial suspension was incubated with equal volume of 200 μg/mL of streptavidin (Thermo Fisher) for 30 min at room temperature on a tube rotator. Excess streptavidin was washed out three times by pelleting nanovials at 2,000 × g for 30 s on a Galaxy MiniStar centrifuge (VWR), removing supernatant and adding 1 mL of fresh Washing Buffer.

**Anti-CD45 and cytokine capture antibody labeled nanovials.** Streptavidin-coated nanovials were reconstituted at a five times dilution in Washing Buffer containing 140 nM (20 μg/mL) of each biotinylated antibody or cocktail of antibodies: anti-CD45 (Biolegend, 368534) and anti-IFN-γ (R&D Systems, BAF285), anti-human NYESO1/HLA-A2 (3M4E5). Nanovials were incubated with antibodies for 30 min at room temperature on a rotator and washed three times as described above. Nanovials were resuspended at a five times dilution in Washing Buffer or culture medium prior to each experiment.

**pMHC labeled nanovials.** MHC monomers with peptides of interest were synthesized and prepared according to a published protocol^70^. Streptavidin-coated nanovials were reconstituted at a five times dilution in Washing Buffer containing 20 μg/mL biotinylated pMHC and 20 µg/mL biotinylated anti-human IFN-γ antibody.

